# Structure of a new capsid form and comparison with A-, B- and C-capsids clarify herpesvirus assembly

**DOI:** 10.1101/2025.03.19.644230

**Authors:** Alexander Stevens, Saarang Kashyap, Ethan Crofut, Ana Lucia Alverez-Cabrera, Jonathan Jih, Yun-Tao Liu, Z. Hong Zhou

## Abstract

Three capsid types have been recognized from the nuclei of herpesvirus-infected cells: empty A-capsids, scaffolding-containing B-capsids, and DNA-filled C-capsids. Despite progress in determining atomic structures of these capsids and extracellular virions in recent years, debate persists concerning the origins and temporal relationships among these capsids during capsid assembly and genome packaging. Here, we have imaged over 300,000 capsids of herpes simplex virus type 1 by cryogenic electron microscopy (cryoEM) and exhaustively classified them to characterize the structural heterogeneity of the DNA-translocating portal complex and their functional states. The resultant atomic structures reveal not only the expected A-, B-, and C-capsids, but also capsids with portal vertices similar to C-capsids but no resolvable genome in the capsid lumen, which we named D-capsids. The dodecameric dsDNA-translocating portal complex varies across these capsid types in their radial positions in icosahedral capsids and exhibits structural dynamics within each capsid type. In D-capsids, terminal DNA density exists in multiple conformations including one reminiscent to that in C-capsids, suggesting D-capsids are products of failed DNA retention. This interpretation is supported by varying amounts of DNA outside individual D-capsids and by correlation of capsid counts observed in situ of infected cell nuclei and those after purification. Additionally, an “anchoring” segment of the scaffold protein is resolved interacting with the portal baskets of A- and B-capsids but not D- and C-capsids. Taken together, our data indicate that A-capsids arise from failed DNA packaging and D-capsids from failed genome retention, clarifying the origins of empty capsids in herpesvirus assembly.

**Significance:** As the prototypical herpesvirus, herpes simplex virus 1 (HSV-1) exhibits a global seroprevalence of 67% and approaching 90% in some localities. Herpesvirus infections can cause devastating cancers and birth defects, with HSV-1 infections leading to cold sores among the general population worldwide and blindness in developing nations. Here, we present atomic structures of the capsids sorted out from the nuclear isolates of HSV-1 infected cells, including the previously recognized A-, B-, and C-capsids, as well as the newly identified D-capsid. The structures show the details of protein-protein and protein-DNA interactions within each capsid type and the positional and interactional variability of the viral DNA-translocating portal vertex among these capsids. Importantly, our findings suggest that A-capsids are products of failed dsDNA packaging and D-capsids of failed genome retention. Together, the high-resolution 3D structures clarify the processes of genome packaging, maintenance, and ejection during capsid assembly, which are conserved across all herpesviruses.

## Introduction

First described by the Greek historian Herodotus over 2000 years ago, human herpes simplex virus 1 (HSV-1) is a highly adapted linear double-stranded DNA (dsDNA) virus which evolved alongside humans since our last common ancestor (1, 2). HSV-1 is a prototypical member of the *Herpesviridae* family and infects two-thirds of humans worldwide, with regional seroprevalence as high as 90% (3). Infection typically presents as mild symptoms in healthy individuals, such as cold sores in the orofacial region, but can lead to serious disease amongst the immunocompromised or immune naïve. Indeed, HSV-1 is the leading cause of lethal encephalitis in the United States and a major cause of blindness worldwide, and the lack of efficacious treatments or prophylactics makes HSV-1 a serious global health challenge (4, 5).

Ever since the first 30-40 Å resolution 3D reconstructions of HSV-1 capsids some 35 years ago (6, 7), significant progress has been made in determining and understanding the virion structure and genome packaging (8–10). HSV-1 packages its 152 kbp genome into an icosahedral T=16 nucleocapsid 125 nm in diameter, surrounded by a pleomorphic tegument layer and glycoprotein-decorated lipid envelope. The HSV-1 nucleocapsid is composed of 955 copies of the major capsid protein (MCP) arranged into 150 hexons and 11 pentons that decorate icosahedral 5-fold (I5) vertices. A unique portal vertex exists at a twelfth I5 vertex through which the genome is translocated during assembly. 320 triplex protein complexes (Tri), each composed of one Tri1 and two Tri2 (Tri2A & Tri2B) subunits rivet MCP capsomers (i.e. hexons and pentons) together, while small capsid proteins (SCP) decorate the top of each MCP tower. Capsid vertex specific components (CVSC) buttress pentonal MCPs and their neighboring Tri complexes and stabilize the capsid (10).

During infection, HSV-1’s lipid envelope fuses with the host cell membrane and releases tegument proteins and the nucleocapsid into the cytosol (11, 12). The nucleocapsid travels along microtubules and docks at a nuclear pore complex, where the viral genome ejects into the nucleus for transcription (13–16). Nascent structural proteins including Tri1, Tri2, MCP, SCP, scaffolding protein (pUL26.5), protease-containing scaffolding protein (pUL26), and portal protein (pUL6) are then translated in the cytosol and trafficked back to the nucleus. Here, portal proteins assemble into a dodecameric basket-like structure that is thought to seed the assembly of scaffolding proteins, in turn driving the formation of spherical procapsids composed of MCP, SCP, and triplexes (17–22). A separate protein complex, the terminase, associates with the dodecameric portal and initiates genome packaging and the concurrent autoproteolytic cleavage of the scaffold (23–27). Subsequent capsid maturation steps result in a range of outcomes: successful genome packaging produces C-capsids, while aberrant byproducts also form, like B-capsids that have angularized but failed to release scaffolding (28) and A-capsids that have released scaffolding but generally do not contain visible genome. Only C-capsids ultimately become infectious virions. (29). A-, B-, and C-capsids are readily distinguishable in singular electron micrographs by their capsid lumens, as C-capsids have a fingerprint-like appearance created by the genome, B-capsids have an obvious scaffolding core, and A-capsids appear largely empty. Even with a high degree of speciation amongst different herpesviruses subfamilies (30–32), capsid assembly leading to the formation of A-, B-, and C-capsids within the host nucleus is largely conserved.

Despite the progress outlined above, debate persists concerning temporal origins and relationships amongst the various capsid outcomes, reflecting a lack of granular understanding of the capsid assembly pathway. And while previous structural studies of DNA-devoid capsids from herpes simplex virus type 2 (HSV-2), varicella zoster virus (VZV), and human cytomegalovirus (HCMV) have shown additional characteristics specific to different capsid types—for example, that A- and B-capsid portals sit lower within the capsid vertex than the “elevated” C-capsid portals (33–36)—little is known about the underlying conformational changes and protein-protein interactions that contribute to such differences. Here, we used extensive computational sorting with an unprecedentedly large dataset of capsid images to resolve the first high-resolution asymmetric reconstructions of the HSV-1 A-, B-, and C-capsids derived from the nuclei of infected cells. In addition to features consistent with previous herpesvirus structures (33, 34, 37), our dataset reveals a previously unidentified capsid end product. These new capsids feature C-capsid-like “elevated” portals and, in many cases, terminal genome within the portal translocation channel, but no organized genome in the capsid lumen. We refer to these capsids as “D-capsids” because their C-capsid-like portal suggests they were originally C-capsids, but the absence of most of the genome indicates that they are degraded. Meanwhile, in high-resolution reconstructions of “lowered” A- and B-capsid portal baskets, we identified densities corresponding to scaffolding proteins where they bind to the portal. Additionally, we resolved the global arrangement of the capsid-bound fragment of the scaffolding protein attached to the inner walls of B-capsids. These results provide novel insights into capsid assembly, genome maintenance, and viral maturation.

## Results

### 3D classification of capsids from host nuclei and discovery of new D-capsids

We isolated HSV-1 capsids from the nuclei of infected Vero cells and subjected them to single-particle analysis using cryoEM to determine their 3D structures. We recorded 33,399 movies and picked 309,503 particles. We extracted the portal vertices as sub-particles, which were classified into 30,298 A-, 66,591 B-, and 153,080 C-capsid vertex particles and refined by imposing C5 symmetry to 3.8, 3.7, and 3.5 Å reconstructions, respectively (Fig. S1). Using an established symmetry expansion and relaxation computational procedure (10), we resolved the symmetry mismatch between the C5 symmetrical vertices and C12 symmetrical portal baskets to generate C1 reconstructions at 4.3, 4.0, and 4.1 Å for A-, B-, and C-capsid portal vertices, respectively. These orientations were used to generate asymmetric whole-capsid reconstructions (Fig. 1A through D) for comparison with a previously resolved virion structure (Fig. 1E) (10). To improve the resolution of the portal baskets, we carried out separate sub-particle reconstructions and refined them while imposing C12 symmetry, leading to portal basket structures at resolutions of 3.6, 3.5, and 3.7 Å for the A-, B-, and C-capsids, respectively (Fig. S1).

**Figure 1:**
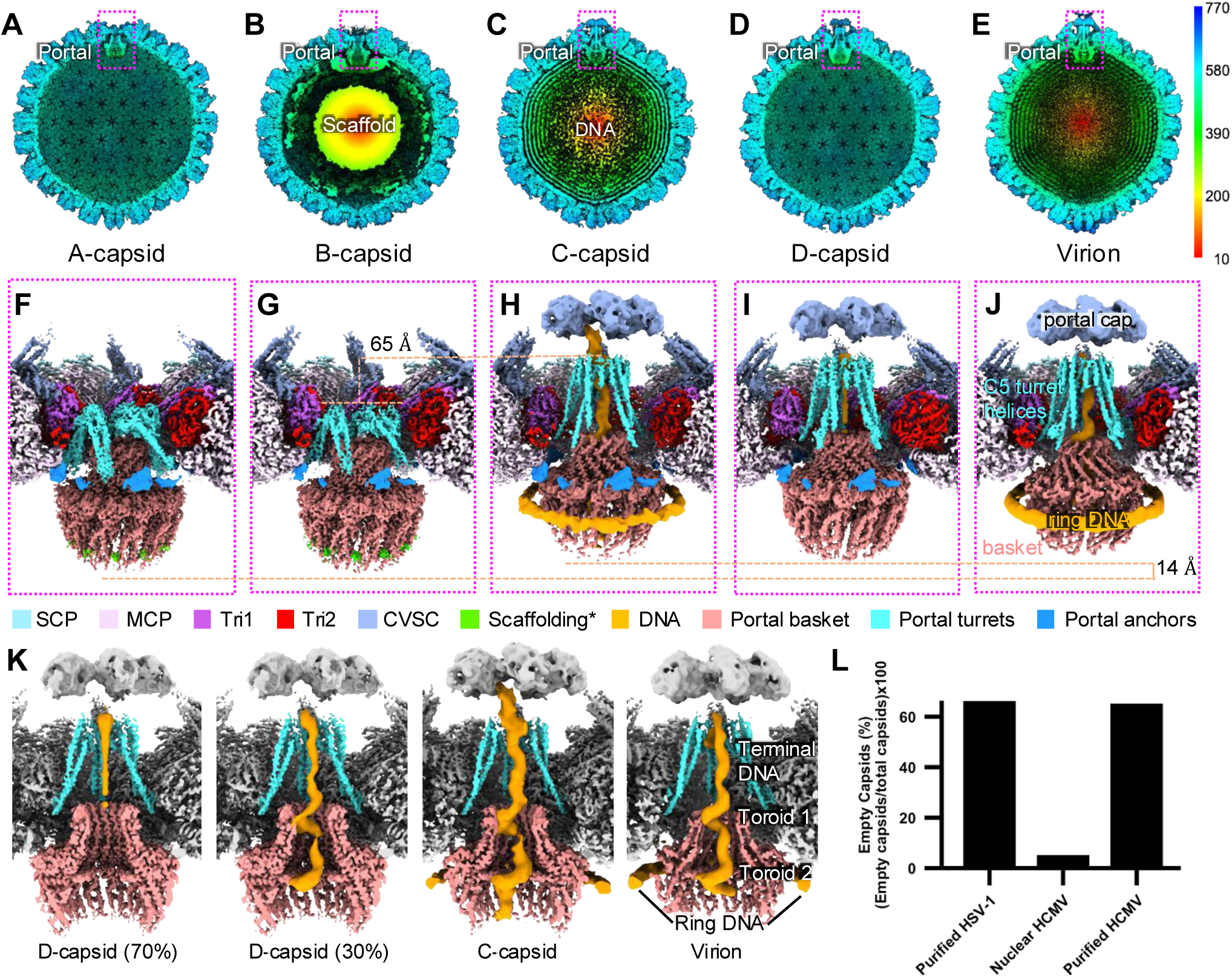
Asymmetric reconstructions and comparisons of HSV-1 A-, B-, C-, D-, and virion nucleocapsids. (A)-(E) asymmetric reconstructions of A-, B-, C-, D-, and virion (EMDB: 9864) capsids showing empty (A,D), scaffold-containing (B), and genome-containing (C,E) capsid shells. The color bar on the right indicates radial distance from the capsid center in Å. (F)-(J) asymmetric reconstructions of the portal vertices from A-, B-, C-, D-, and virion (EMDB: 9860, 9862) capsids in A-E. The orange dashed line in G and H labeled with “65 Å” indicates difference in turret helix height across capsid types. Lower orange dashed line labeled with “14 Å” indicates the difference in portal basket height across capsid types. *Scaffolding density (lime) in F and G are not necessarily equivalent, as the density in A-capsids is probably a fragmentary remnant after proteolytic cleavage. SCP: small capsid protein. MCP: major capsid protein. Tri1: triplex 1 protein. Tri2: triplex 2 protein. CVSC: capsid vertex specific component. (K) Comparison of genome organization in D-, C-, and virion portal vertices, showing consistent arrangement in virions, C-capsids, and some D-capsids, but symmetric, round density in another class of D-capsids. (L) graph of prevalence of empty capsids in our study of HSV-1, as well as nuclear thin-section versus purified samples for HCMV using data from (40).

Classification of the presumed C-capsid portals revealed that 88% of particles lacked both capsid luminal DNA and ring DNA (Fig. 1K and S1). While the absence of ordered luminal genome conflicts with the canonical definition of C-capsids, these unusual portals had been grouped with C-capsids during classification due to having extended turret helices, an elevated portal basket, a portal cap, and genome density inside the DNA-translocating channel of the portal, in contrast to A- and B-capsids where the turret helices are retracted, the portal basket is lower, the cap is not present, and genome density is lacking inside the portal channel (Fig. 1F through J) (10, 38, 39). Surprised by this result, we manually curated whole C-capsid particles, revealing that 135,916 of 153,080 capsid particles appeared in micrographs as partially or totally devoid of luminal genome—like A-capsids—yet still contributed to reconstructions with C-like portal features (Fig. 1I, 1K, S1, and S2). Symmetry relaxation of these C-like portals revealed that ∼30% of particles had toroidal genome organization in the portal channel equivalent to the genome structure in virions and C-capsids, with the other ∼70% harboring less distinct DNA density in the portal channel, perhaps due to variable arrangements (Fig. 1I and K).

These observations led us to investigate the origins of these unusual capsids with C-like portal vertices. Previous studies have suggested that herpesvirus C-capsids isolated from nuclei are prone to degradation (29, 33), so we wondered whether capsids with C-like portals may represent former C-capsids in a degraded state. Indeed, we see varying levels of genome occupation in the micrographs of capsids with C-like portals, often with genome egressing from the capsid shell (Fig. S2). The total proportion of capsids with empty lumens in our study is 66%, which is much higher than the proportion usually observed within cell nuclei infected with herpesviruses (Fig. 1L) (40). This high proportion of empty capsids resembles that of HCMV, which is known to have unstable C-capsids resulting in a higher empty capsid proportion specifically in purified samples (Fig. 1L) (29, 34, 40). These findings suggest that capsids with C-like portals are aberrant products that arise when C-capsids fail to retain viral DNA, and we hereafter refer to these as degraded (D-) capsids.

### Atomic structures of portal complexes and their contacts with the capsid shell

Our large dataset allowed us to generate reconstructions of the portal basket for all nucleus-derived capsid types at far higher resolution than previous herpesvirus structures, enabling modeling of the portal protein’s basket (a.a. 26-307 and 494-623) and turret helices (a.a. 337-473) (10, 33, 35). The basket is a 12-fold symmetrical complex consisting of five domains, including the wing (a.a. 26-62, 150-174, 222-370), stem (a.a. 271-297, 517-541), clip (a.a. 298-307, 494-516), β-hairpin (a.a. 542-557) and crown (a.a. 62-149, 175-221, 558-623) (Fig. 2A). Ten of the twelve monomers from the basket are linked to the turret helices, each of which consists of one long helix extending towards the portal cap and connecting to two smaller helices arranged in a small helical bundle. The two non-contributing portal monomers are presumably disordered or flexible beginning at their turret sequences (Fig. 2A). In C-capsids, D-capsids, and virions, the turret helices are fully extended, whereas in A- and B-capsids, they are retracted and positioned ∼65Å lower (Fig. 1F through J). These features are consistent with previous herpesvirus structures (35, 41, 42).

**Figure 2:**
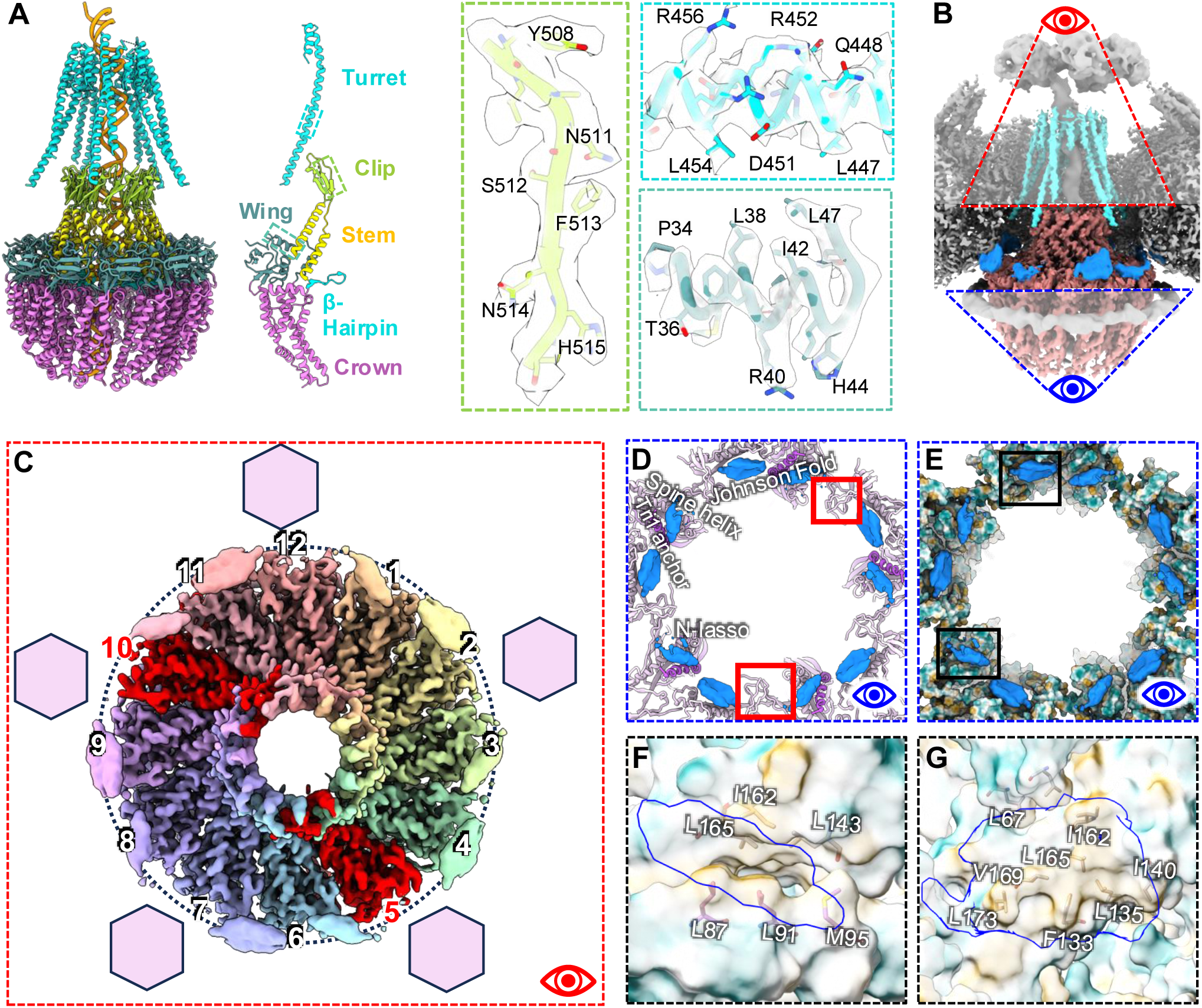
High resolution structure of the portal basket. (A) from left to right: full portal dodecamer including turret helices colored by domain, resolved portions of a portal monomer, and examples of model fit to cryoEM density in the clip (green), turret (cyan), and wing (blue green) domains. (B) cryoEM density of the portal vertex showing orientation of top (red) view in panel C and bottom (blue) view in panels D-E. (C) Top view of the portal basket showing assignment of portal anchors (smooth outer blobs) to portal subunits, with subunits missing an anchor labeled in red. Hexagons indicate fivefold locations of hexons relative to the portal. (D) bottom view of MCP (lavender) and Tri1 (purple) fragments near the portal anchors (dodger blue) with important domains of MCP and Tri1 labeled. (E) bottom view of MCP and Tri1 fragments near portal anchors colored by hydrophobicity. Bottom left and top black outlines indicate the views in panels F and G respectively. (F)-(G) views of the lone helix (left) and helix pair (right) of the portal anchors (blue outlines) overlayed with hydrophobic surface view of their binding pockets, with hydrophobic residues shown.

In addition to the basket and turrets, the portals of all capsid types feature 10 helical densities—five helix pairs and five lone helices—originating from the N-terminal wing domains of 10 of the 12 portal subunits (Figs. 1F through I and 2B through C). These appear to anchor the portal basket to the capsid shell (Fig. 2B and C), forming an arrangement similar to the “10-helix anchor” seen in HCMV (Li et al., 2023). The helix pairs interface with hydrophobic pockets defined by MCP Johnson-folds and dimerization domains adjacent to the portal (a.a. 143-145, 165-176, 342-346), while the lone helices interface with the leucine-rich Tri1 N-terminal anchors (a.a. 82-92) and MCP Johnson-fold domains (a.a. 160-170) (Fig. 2D through 2G). When viewed from outside the capsid, two of the twelve portal subunits corresponding to the 5 and 10 o’clock positions did not have N-terminal anchors (Fig. 2C). This 12-fold to 5-fold symmetry mismatch is similar to that observed in the turret helices, where two portal subunits are conspicuously absent from the turret arrangement.

### Portal basket positioning and flexibility varies across capsid types

The flexibility of the portal basket is essential to accommodate the structural changes and mechanical demands associated with genome packaging, ejection, and the transition between capsid states. To characterize the portal’s structural variability, we used the 3D flexible refinement (3DFlex) tool from cryoSPARC to model the movement of the portal basket relative to the capsid shell (43). All capsid types showed some variability in portal basket position and rotation relative to the capsid shell. A- and B-capsids varied in portal height by ∼2 and ∼1 nm respectively, could rotate up to 13 degrees relative to the capsid shell, and exhibited up to 1 nm of portal basket “sway”, due to flexibility between the clip and nearby wing and stem domains (Fig. 3A, 3B, and Movie S1 through 4). C-capsid portals did not vary in height but were still capable of 9-degree rotations and ∼1 nm of portal basket sway (Fig. 3C and Movie S5 though 6). D-capsid portals had 4 Å variability in height but otherwise resembled C-capsid portals (Fig. 3D and Movie S7 through 8). These findings reveal distinct patterns of portal basket variability, with A- and B-capsids exhibiting greater positional and rotational flexibility compared to the more constrained movement observed in C- and D-capsids.

**Figure 3:**
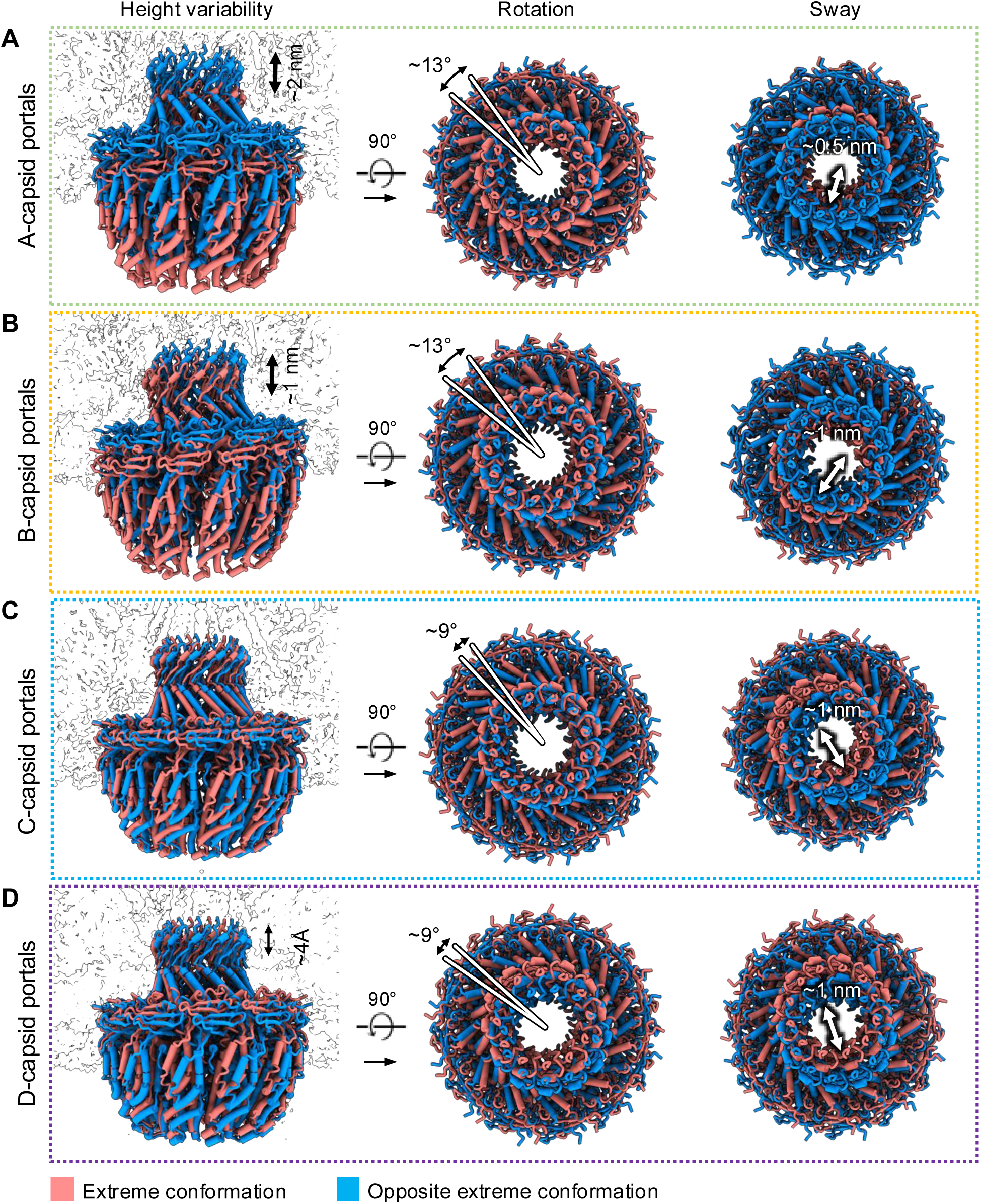
Portal basket flexibility and positional variability within each capsid type for HSV-1 A-, B-, C-, and D-capsids. (A) – (D) Portal basket models fit into alternative conformations found via 3D flex analysis for each of the four capsid types,with the extreme edges of their conformational ranges represented as salmon and aqua structures. Multiple separate analyses show vertical translation (left), rotation (center), and a swaying motion of the top half of basket (right).

### Scaffolding protein is anchored to A- and B-capsid portal baskets and B-capsid shells via conserved hydrophobic motifs

The C12 reconstruction of both A- and B-capsid portal baskets revealed a small hook-like density with large aromatic side chains and sharp turns wedged into a hydrophobic pocket between the crown domains of each portal monomer (Fig 4A through B and 4F through G). This density is weaker in A-capsids, but still conspicuous, whereas it is completely absent in C-capsids, D-capsids, and mature virions. (Fig. 4C through E, 4H through J). It neighbors W90 and W127 in the portal monomers, known to be essential for portal-scaffold interactions (44). Investigators previously showed that portal protein residues 449-457—particularly Y451, P452, and E454 (44, 45)—are essential to the portal-scaffold interaction and thus efficient capsid assembly (44). Indeed, we found the scaffolding protein residues 449-455 (PYYPGEA, numbered relative to pUL26) to be a good fit for the small density (Fig. 4K and L). Based on our model, P452 interacts with the essential residues W127 and L131 (both adjacent to W90) from the first portal monomer, whereas Y451 forms a hydrophobic interface with Y83 and Y608 from the second portal monomer (Fig. 4L and M). Further, E454 from the scaffolding appears to stabilize Y608 via hydrogen bonding, buttressing the hydrophobic interactions between the three aromatic residues (Fig. 4L). Sequence alignment of scaffolding and portal proteins showed that Y608 and Y83 are highly conserved across herpesviruses, suggesting this motif is essential for portal-scaffold association (Fig. 4N).

**Figure 4:**
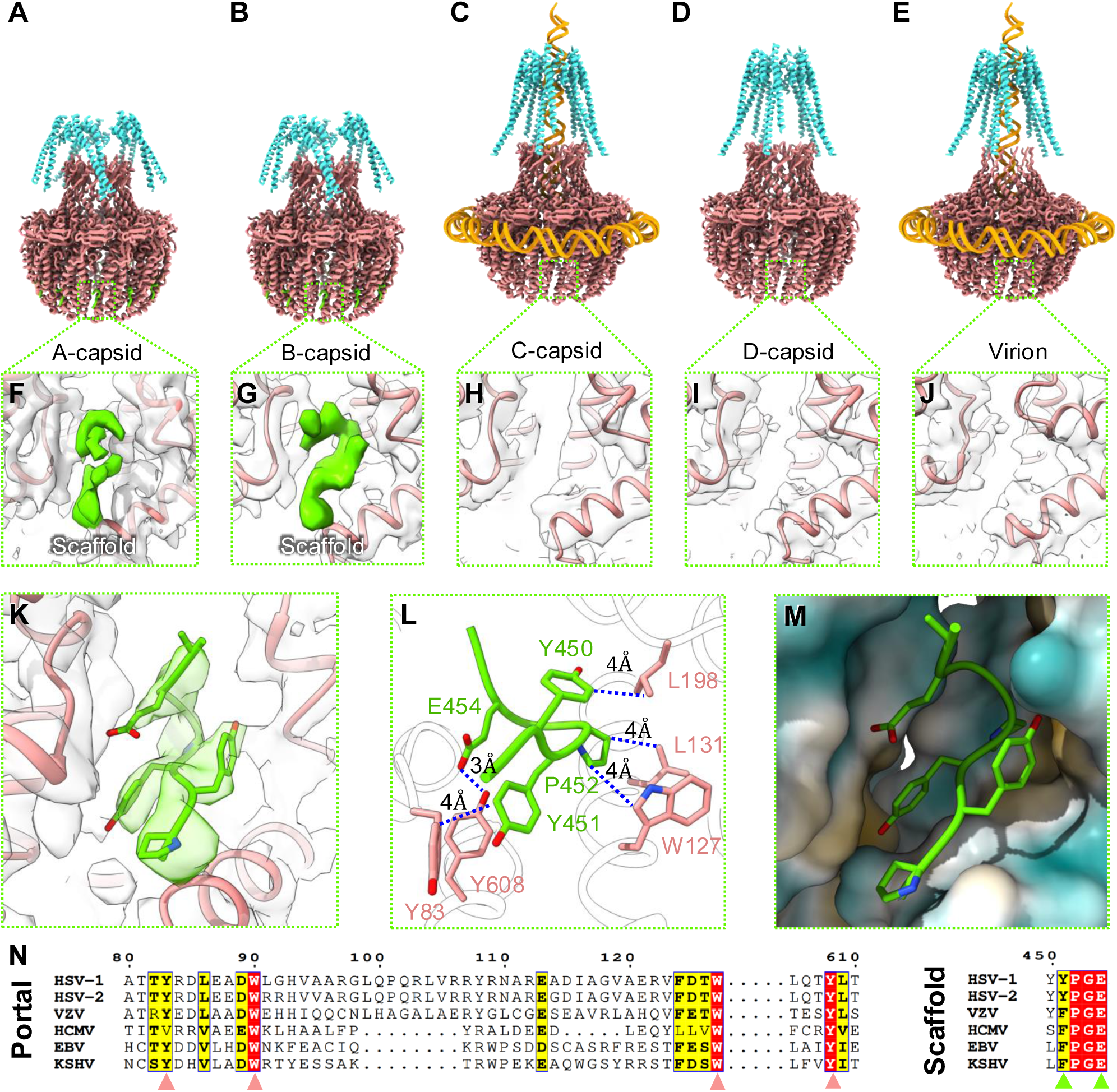
The scaffold protein interacts with the portal dodecamer. (A)-(E) ribbon diagram of modeled residues in the portal dodecamer for A-, B-, C-, D-, and virion capsids, including basket (salmon), turret helices (cyan), and where applicable, the associated genome (orange) and scaffolding protein loops (lime). (F)-(J) zoom-ins of the ribbon model and density of the portal’s scaffold binding pocket for the five capsid types in (A)-(E), showing presence or absence of scaffold density (lime). (K) Residues 449-455 of the scaffolding protein fit into a density interacting with the portal. (L) Interatomic distances between atoms of scaffolding and portal at their interaction site, highlighting hydrophobic residues. (M) Scaffolding loop shown with a hydrophobicity map of the portal, showing hydrophobic spots near Y451 and P452. (N) Sequence alignment of portal and scaffolding proteins across a range of herpesviruses, demonstrating sequence conservation of residues important for portal-scaffold interactions.

Despite the high-resolution details revealed in residues 449-455 of the scaffolding protein, the rest of the scaffolding complex remained poorly resolved both around the portal basket and capsid lumen (Fig. 1B, and 5A through C). Our asymmetric reconstruction of B-capsids shows that the portal basket is encircled by large prominent densities (Fig. 1B), which we attribute to scaffolding protein that remains attached to the capsid shell (Fig. 5A). Further, these densities, and others similar, appear to originate from local 3-fold axes near the positions of proximal Tri1 N-terminal anchors (Fig. 5A), consistent with observations of the scaffolding anchor domain in HCMV (34) (Fig. 5C). These densities are noticeably absent underneath the local 3-fold axes immediately adjacent to pentons (Fig. 5A through 5C).

**Figure 5:**
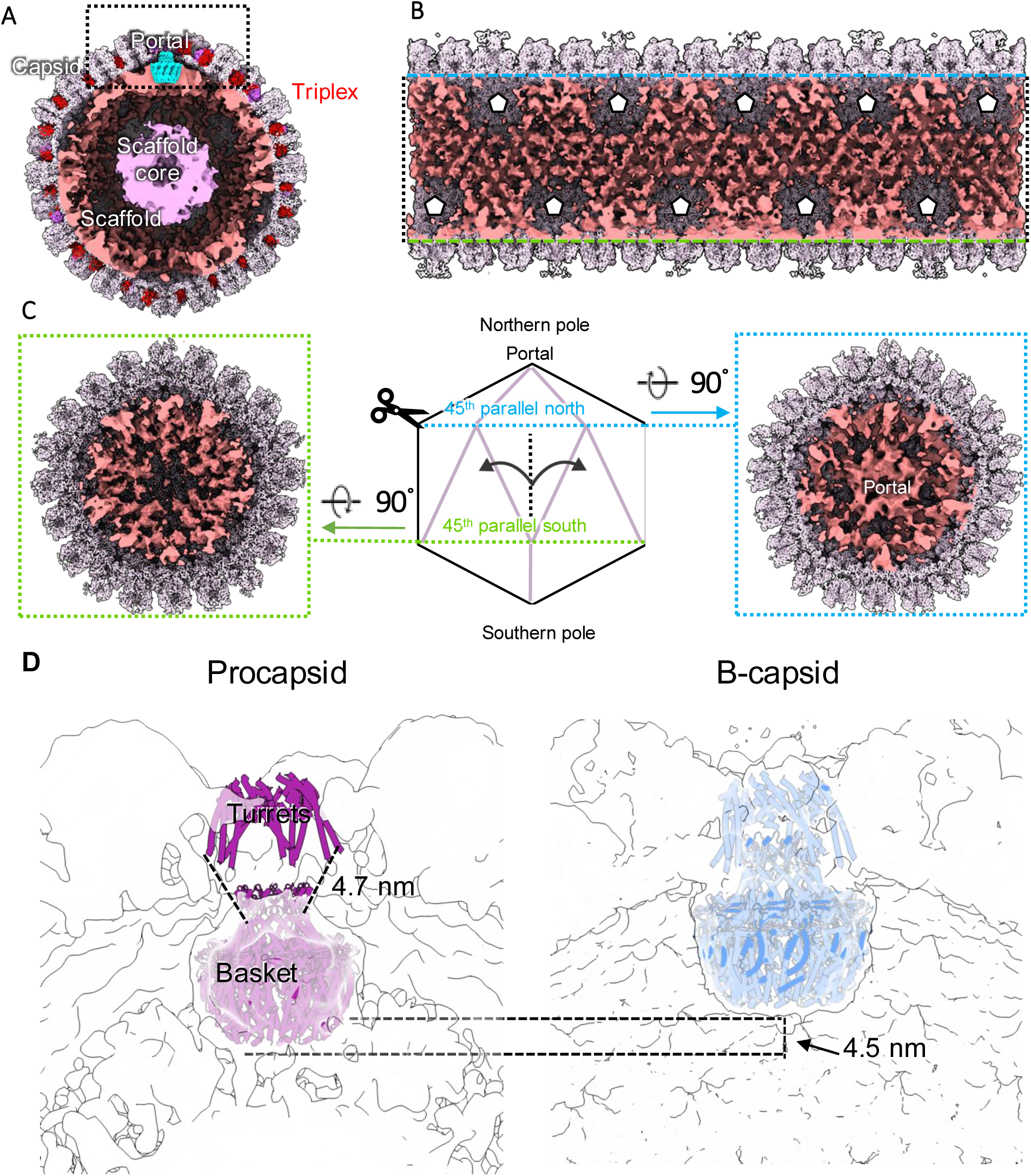
Global arrangement of the scaffold in whole B-capsids. (A) Cut view of whole B-capsid reconstruction showing large internal scaffold densities (top left), with Tri1 and Tri2 labeled in purple and red, respectively. (B) Unwrapped view of the capsid floor showing global scaffold arrangement, with penton locations marked with black-outlined white pentagons. (C) Scaffold densities at portal (blue dashed outline) and antipodal (green dashed outline) poles. (D) Comparison of the portal basket in HSV-1 procapsids from Buch et al. with our B-capsid reconstruction, showing 4.5 nm difference in position.

## Discussion

Characterizing herpesvirus capsid assembly requires thorough sampling to capture diverse intermediates and end products in viral replication. Here, we used single-particle cryoEM to resolve the first near-atomic resolution asymmetric structures of HSV-1 capsids derived from the nuclei of infected cells. Our large dataset enabled the identification of a new capsid form, the D-capsids, and provided the highest resolution detail of a herpesvirus portal to date, including its interactions with the scaffold and dynamics with the capsid shell. These findings shed light on longstanding questions about how the genome impacts portal basket position, the portal’s role in procapsid formation, the portal-scaffold interaction, and the origins of empty capsids.

The presence or absence of genome within the portal translocation channel appears to be related to the position of the portal basket. C-capsid portal baskets are positioned higher at the portal vertex compared to their A- and B-capsid counterparts, which has been attributed to the densely packed genome pushing the portal up against the capsid shell (34). However, our D-capsid portal structures have the same portal basket position as C-capsids, despite being mostly devoid of genome (Fig. 1K). These observations together suggest that genome pressure alone does not determine portal basket height, and the elevated portal position observed in C- and D-capsids may instead be an irreversible step during capsid maturation.

The portal is important for both procapsid assembly and genome packaging, but the structural basis enabling these dual roles had not been previously characterized. Our analyses of portal basket variability revealed that non-mature A- and B-capsid portals permit more rotational flexibility than mature-state C-capsid portals, which may be important for the dynamic process of genome packaging, during which torque is applied to the genome substrate (34, 46, 47). Additionally, although A- and B-capsid portal baskets can vary in elevation by up to ∼2 nm, they cannot descend to the position of the procapsid portal basket, which is further situated 4.5 nm lower with respect to the capsid shell (Fig. 5D) (48). Indeed, the linker residues connecting the A- and B-capsid portal baskets to their capsid-docked N-terminal anchors are too short to permit such a 4.5-nm shift, suggesting that the procapsid basket must not be initially attached to the capsid shell via N-terminal anchors. This suggests that during angularization, the portal basket moves away from the scaffold core, while its N-terminal anchors attach irreversibly to the capsid shell, limiting the basket’s positional variability and preventing it from descending back to its procapsid position. Altogether, the ability of the portal basket to begin in a lowered conformation in procapsids may be critical for proper interaction with the scaffold and thus the nucleation of capsid assembly, while the subsequent attachment of portal N-terminal anchors post-angularization restrains the basket near the capsid shell, enabling it to sustain robust terminase-driven forces during genome packaging.

Our structures unveil the scaffolding proteins’ interactions with the portal and capsid shell, which were previously suggested to be important for viral assembly (44, 49–51) but lacked structural detail. Herpesvirus procapsid assembly is promoted by scaffold-scaffold interactions, enabling the progressive addition of scaffold-capsid complexes into partially formed capsids (19). Our B-capsid structure shows scaffolding densities emanating from under local 3-fold axes in close proximity to Tri1 N-anchors, consistent with observations in protease-null mutants (52). While their exact interface to the B-capsid luminal wall remains unclear, the binding pocket harboring the Tri1 N-anchor is known to be promiscuous, variably binding the portal N-anchor in our study and in HCMV (34) as well as isomeric forms of MCP Johnson-fold components at penton locations (9, 10). This raises the possibility that during early procapsid assembly stages, these pockets may bind scaffold protein C-terminus, which may later be displaced by Tri1 N-terminal anchors after scaffolding protease activation (34, 52, 53). In addition to scaffold-capsid interactions, our structure shows high-resolution details of the scaffold-portal interface, enabling the first atomic model of a portion of the scaffolding protein and revealing its tightly packed, hydrophobic nature nestled between portal monomers (Fig. 4F, G, and K through M). This finding is likely relevant in other herpesviruses, as the relevant residues on the scaffolding and portal proteins are conserved (Fig. 4N), in addition to direct structural evidence of a portal-scaffold interaction at an equivalent motif in HCMV (34). Thus, this interface may serve as an attractive target for the development of pan-herpesvirus inhibitors of capsid assembly.

Prior to the current study, A-capsids were thought to result from unsuccessful genome packaging, but their exact origins were uncertain. Our A-capsid portal reconstructions still contain scaffolding hook densities, although with weaker density than in B-capsids, suggesting low occupancy, but even residual presence of the scaffolding protein suggests that these capsids were likely never fully packaged with genome, as full genome content in the capsid lumen should displace the portal-bound scaffold. However, because D-capsids generally appear with empty lumens in micrographs, it is likely that D-capsids have been observed in past studies of herpesviruses but were grouped with A-capsids. Therefore, the “empty” capsids may be formed at various stages of assembly, both before and after the completion of genome packaging for A- and D-capsids respectively (Fig. 6), providing partial validity to the A-capsid multi-origin model put forward in Kaposi’s sarcoma-associated herpesviruses (54).

**Figure 6:**
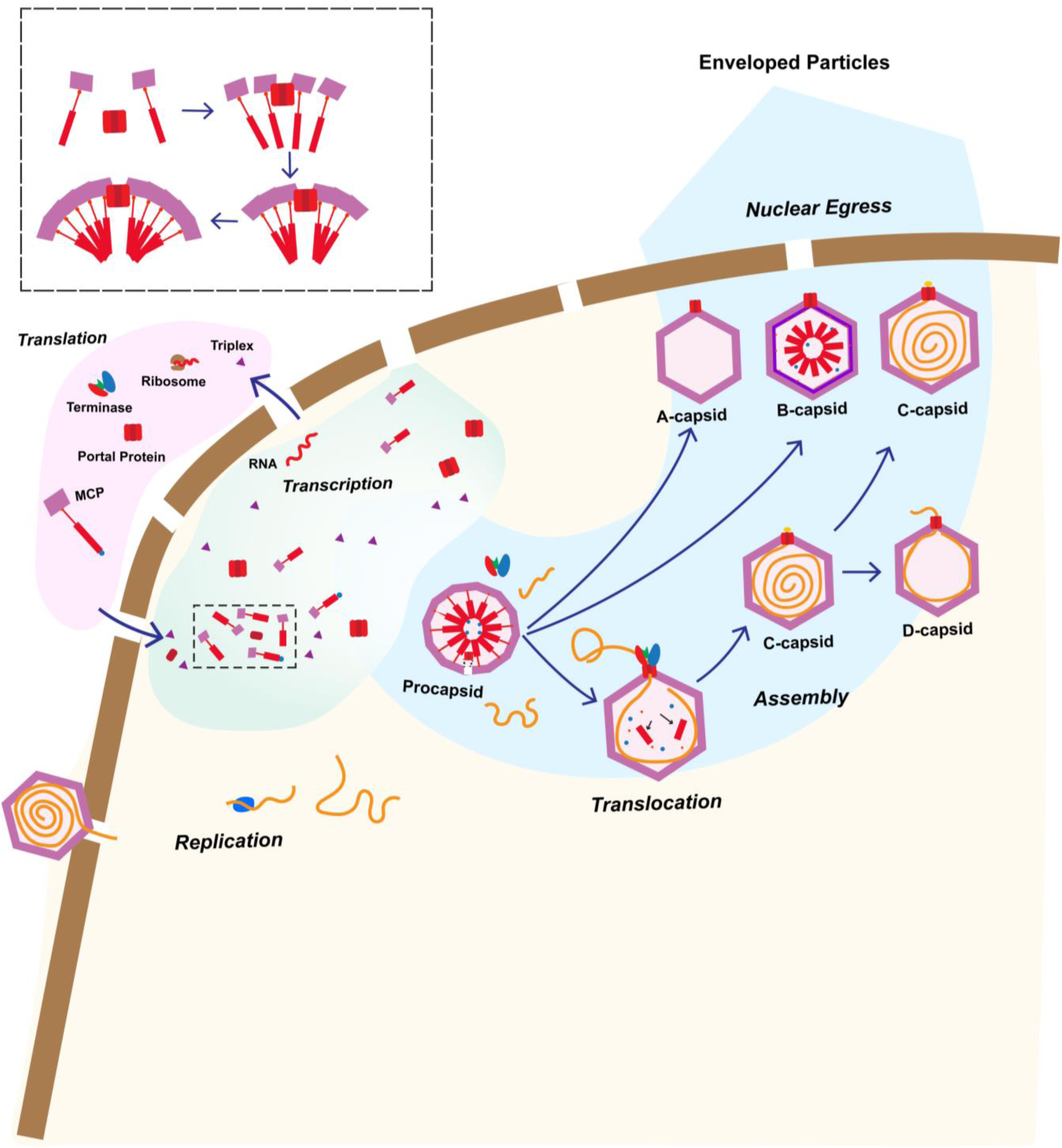
Proposed model of herpesvirus maturation and assembly. Virion capsids release the viral genome into the host cell nucleoplasm, where the genome is transcribed and translated to form viral proteins. These proteins are then imported into the nucleus and associate with a free portal complex via scaffold-mediated interactions, initiating assembly of the procapsid. If angularization occurs prematurely, B-capsids are formed. If angularization and scaffold digestion are completed but not genome packaging, A-capsids are formed. Successfully packaged capsids, or C-capsids, may either undergo further maturation into virions or fail to retain genome, resulting in the formation of D-capsids.

In summary, 3D structural classification enabled by an unprecedentedly large dataset of HSV-1 capsid images has led to the discovery of a capsid representing a temporally distinct state compared to previously described capsids, and alongside our high-resolution structures of the portal vertices, provide new insight into herpesvirus capsid assembly. Our structures reveal atomistic details of portal-type specific interactions and more clearly delineate A-, B-, C-, and newly defined D-capsids. The identification of D-capsids as a distinct end product resulting from C-capsid degradation clarifies the nature of observed empty capsids and conformational changes underpinning herpesvirus genome packaging and ejection. A more complete choreography of herpesvirus assembly inside host nucleus (Fig. 6) and their atomic structures also offers a framework for the development of structure-based antiviral therapies targeting portal function.

## Methods

### Virus strain and cell line

The HSV-1 strain KOS (VR-1493; ATCC, Manassas, VA, USA) was used in this study. African green monkey kidney cells (Vero) (CCL-81; ATCC, Manassas, VA, USA) served as host for virus propagation and were maintained in culture media DMEM (Cat No. MT10-017-CV; CORNING, New York, NY, USA) supplemented with 10% heat-inactivated fetal bovine serum (FBS) (Cat No. S11150H; R&D systems, Minneapolis, MN, USA), Penicillin (100 IU/mL) and Streptomycin (100 µg/mL) (Pen/Strep) (Cat No. 30-002-Cl; CORNING, New York, NY, USA), at 37 °C in a humidified atmosphere of 5% carbon dioxide (CO_2_) and 95% oxygen O_2_).

### Preparation of HSV-1 capsids

Vero cells at 90-95% confluence, grown in 18 T175 (175 cm^2^ area) tissue culture flasks, were infected with 0.4 mL of HSV-1 stock dispersed in 38 mL of freshly supplemented cell culture media per flask. After 48 h post-infection the cells were harvested and centrifuged at 10,000 × g for 20 min in a Fiberlite F14-6 × 250y Fixed-Angle Rotor (Thermo Fisher Scientific, Waltham, MA, USA) at 4 °C. All subsequent sample purification steps were carried out at 4 °C. The cell pellet was washed with phosphate buffered saline (PBS) of pH 7.4 (Cat No. 10010049; GIBCO, Carlsbad, CA, USA), and centrifuged at 2,000 × g for 10 min in a SX4750 Swing Bucket Rotor (Beckman Coulter Life Sciences, San Jose, CA, USA). To remove the cell membrane and pelleting the nucleus, the cells were resuspended in 40 mL of membrane lysis buffer (PBS pH 7.4, 0.5 % NP40, 1 X Protease Inhibitor cocktail), kept in ice for 30 min (mixing by gentle inversion of the sample a few times every 10 min) and centrifuged at 2,000 × g for 15 min in a SX4750 swing bucket rotor (Beckman Coulter Life Sciences, San Jose, CA, USA); this step was repeated twice. The nuclear pellet was resuspended in 25 mL of extraction buffer (PBS pH 7.4, 1 mM EDTA, 1X protease inhibitor cocktail), and placed on orbital shaker at low speed for 40 min. Nuclear envelope was lysed via mechanical sheering during 5 consecutive passes of the suspension through a 23-gauge needle. The nuclear lysate was centrifuged at 10,000 × g in a SW 28 Ti swing bucket rotor (Beckman Coulter, Brea, CA, USA) for 1 h, to remove membrane components. The clarified supernatant was placed atop a 30-50% sucrose cushion and centrifuged at 100,000 x g in a SW 28 Ti S swing bucket rotor (Beckman Coulter, Brea, CA, USA) for 1 hr. 1 mL of solution was carefully drawn from the 30-50% sucrose interface, diluted in 10 mL of PBS pH 7.4 with 1 mM EDTA, and pelleted at 100,000 x g in a SW 28 Ti swing bucket rotor (Beckman Coulter, Brea, CA, USA) for 1 hr. The pellet was then carefully dispersed in 15 µL of of PBS pH 7.4 with 1 mM EDTA, preceding to cryoEM grid preparation.

### CryoEM of HSV-1 capsids

HSV-1 capsids were prepared for cryoEM analysis by applying 2.5 µL of sample to 200 mesh holey quantifoil carbon grids which had been glow discharged for 30 seconds. Each grid was flash frozen in liquid ethane using a custom-made plunge-freezing device prior to transfer and storage in liquid nitrogen. CryoEM movies were recorded using a Titan Krios G1 electron microscope (FEI) upgraded with a K3 direct electron detector and Gatan Imaging Filter (GIF) and operated at 300 keV. Movies were recorded in super-resolution mode at a nominal magnification of 81,000 X with a calibrated pixel size of 0.55 Å at the specimen level and estimated electron dose of ∼45 e^-^/ Å^2^. Using SerialEM, 33,399 TIF movies were collected.

### CryoEM image processing

Image alignment, motion correction, and contrast transfer function (CTF) estimation were performed using MotionCor2 and GCTF respectively on 2 X binned micrographs (55, 56). Topaz automated particle picking was performed to pick all 309,503 HSV-1 nucleocapsids within the dose-weighted micrographs, preceding import to RELION (57, 58). To reduce computation time, capsids were boxed and extracted with 1/4 cropping in Fourier space before being subjected to 3-dimensional refinement with icosahedral symmetry enforced. This yielded an initial reconstruction of all capsids at ∼8.8 Å.

We next combined stepwise symmetry expansion and relaxation with sub-particle reconstructions to both improve local resolution and identify the asymmetric portal vertices on each capsid as described previously (10). Briefly, we used RELION’s symmetry expansion utility with I3 symmetry selected to generate 60 equivalent orientations for each capsid particle with different combinations of Euler angles for rotation, tilt and psi. To remove redundant rotations about the Z-axis, which have identical tilt and psi angles we kept only 1 of each set of 5 copies, to yield 12 unique orientations for each particle. Each orientation could then be used to identify the position of one 5-fold vertex in Cartesian coordinates, so that there are 12 in total for each particle which can be reextracted for processing as sub-particles. Portal sub-particles identified for particles whose portals were in frame during movie recording, using exhaustive 3D classification in RELION. This yielded 249,897 portal particles, less than the number of total capsids as some capsids were only partially in frame in the electron micrographs (Fig. S1). Portal particles were then differentiated by their turret helices and separated into “C-like” and “B-like” portal groups. Next the whole particles were manually sorted to identify A-capsids, and these were separated from C-like and B-like portal groups, yielding C-portals, B-portals, B-like A-portals, and C-like portals (D-portals).

Following sorting, sub-particle stacks were transferred to cryoSPARC for further processing (59). Particles were initially subjected to 3D refinement with imposed C5 symmetry, which yielded resolutions of 3.8, 3.7, and 3.5 Å respectively. We then continued with stepwise symmetry expansion and relaxation to resolve each portal types 12-fold symmetrical basket to 3.6, 3.5, 3.7 Å respectively (10). In the case of C-particles we also employed this technique to resolve the C1 reconstruction of the genome segment within the portal.

### Scaffolding density reconstructions

During processing of complete B-capsids, cryoEM map density was observed along the luminal wall of the capsid which appeared to originate from the faces of the icosahedron and beneath the portal vertex. To better resolve these densities, we performed 3D classification of B-portal sub-particles which had been symmetry expanded using the *symmetry expansion* utility in cryoSPARC and a focused classification using a mask which included regions beneath the capsid shell. This yielded large densities which appeared to be protruded regions near the triplex N-terminal anchor and surround the portal baskets in portal sub-particles.

### Model building

Atomic modeling of the HSV-1 peripentonal and portal asymmetric units has been described previously (10, 60). For both assemblies, we modeled previously unresolved loops of MCP, Tri1, Tri2A, and Tri2B combining the *Build Structure* function in ChimeraX and *real-space refinement* in Coot (61, 62). To model the complete asymmetric units, we utilized sub-particle maps from the icosahedral vertices and faces of the viral capsids to assign new helical densities beneath triplexes as the N-terminal anchor of Tri1. We resolved intermolecular clashing in our updated portal and penton asymmetric units using *ISOLDE* from ChimeraX and iterative refinement in Phenix (63).

We flexibly fitted published ab-initio models of the portal dodecamer (10) into our sub-particle portal vertex maps for A-, B-, and C-capsids in *ISOLDE* (64). To model structural flexibility, we fit these updated portal basket models into sub-particle 3D Flex maps of B- and C- baskets.

## Data availability

The atomic models have been deposited in the PDB with accession codes: XXXX. The cryoEM map has been deposited in the EMDB with accession codes: EMD-XXXXX.

## Acknowledgements

This research was supported in part by grants from the National Institutes of Health (R01DE025567 to Z.H.Z and T.-T.W. and R01AI151055 to Z.H.Z.). J.J. acknowledges support from the NIH, National Institute of Arthritis and Musculoskeletal and Skin Diseases grant (NIH 5T32AR071307). E.C. acknowledges support from the Suggs, Ehrisman, Boyer, and Henry endowments, provided by the UCLA Undergraduate Research Center-Sciences. We acknowledge the use of resources in the Electron Imaging Center for Nanomachines supported by UCLA, Core Center of Excellence in Nano Imaging (CNI) at USC, and grants from NIH (S10RR23057, S10OD018111, U24GM116792, and T32GM145388) and NSF (DBI-1338135). We also acknowledge use of resources from the UCLA AIDS Institute, the James B. Pendleton Charitable Trust, and the McCarthy Family Foundation.

## Author contributions

Z.H.Z. conceived the project. A.C.A-C grew and isolated virus, prepared cryoEM samples and obtained cryoEM images; A.S. processed the cryoEM images and determined the structures with assistance from Y.-T.L. A.S., S.K., E.C. and Z.H.Z. interpreted the results and wrote the manuscript; all authors reviewed and approved the manuscript.

---

**Figure S1:**
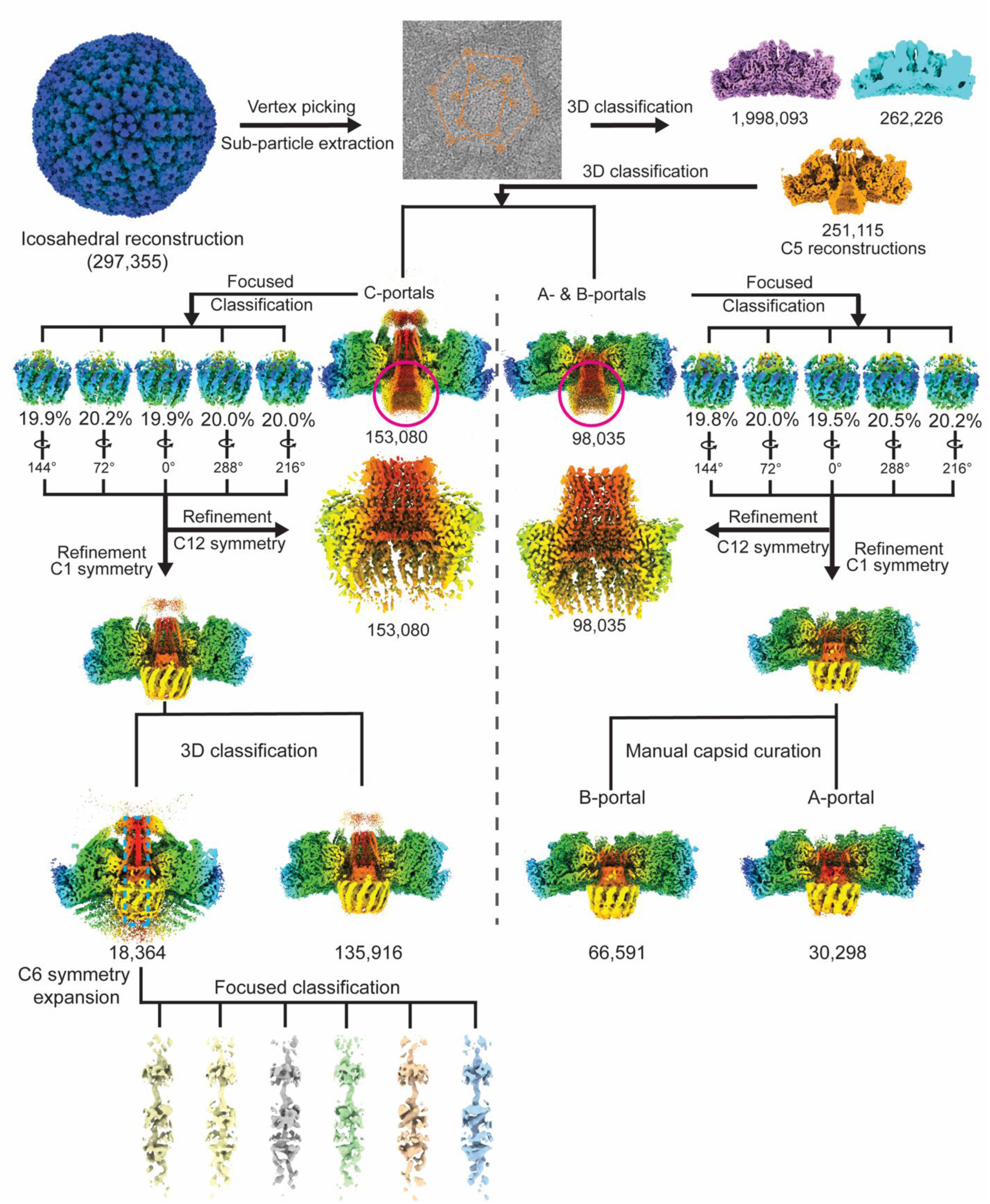
CryoEM image processing workflow for portal vertex reconstructions.

**Figure S2:**
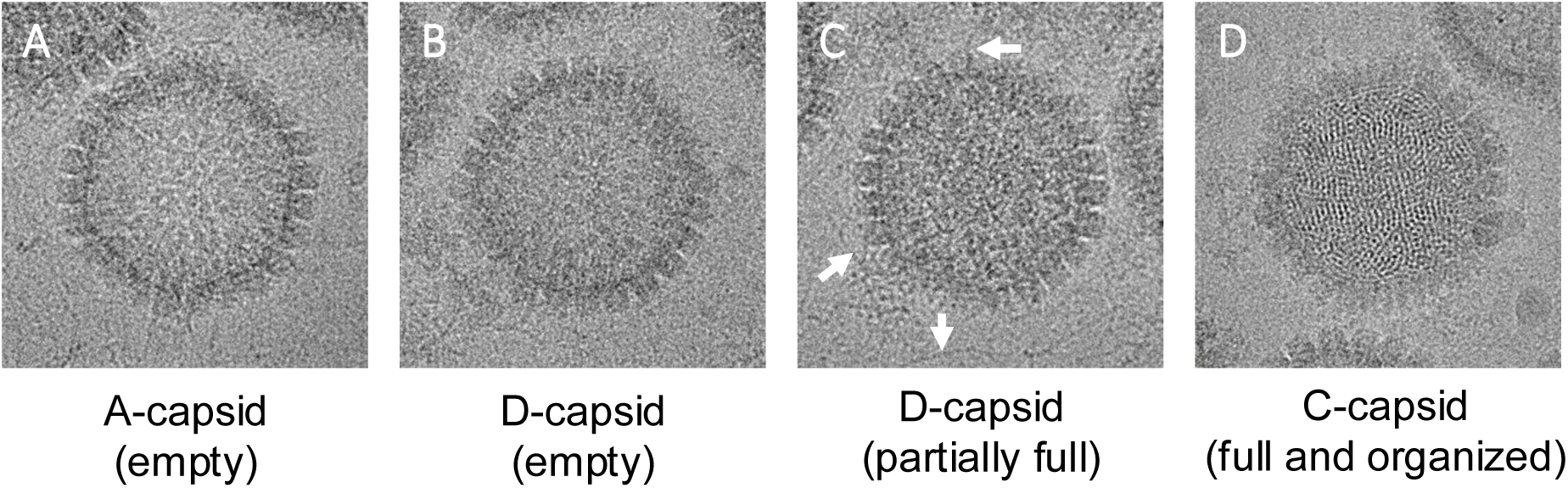
D-capsids occur in a spectrum of genome occupancies. (A) Typical A-capsid micrograph, appearing as an empty capsid. (B) D-capsid micrograph appearing empty, like an A-capsid. (D) Another D-capsid micrograph, appearing filled but not clearly organized. D-capsids occur as a continuum of capsid occupancies, with some empty and some partially filled, but lack the clear genome organization of C-capsids. White arrows indicate free nucleic acid strands, which are common in the sample and may be due to genome ejection in D-capsids. (D) Typical C-capsid micrograph, displaying clear organization of the genome.

**Figure S3:**
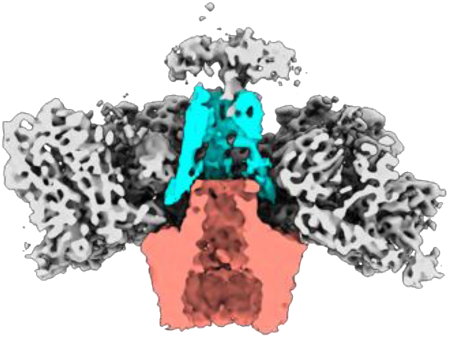
Cross section of D-capsid portal vertex without terminal genome.

**Movie S1.**
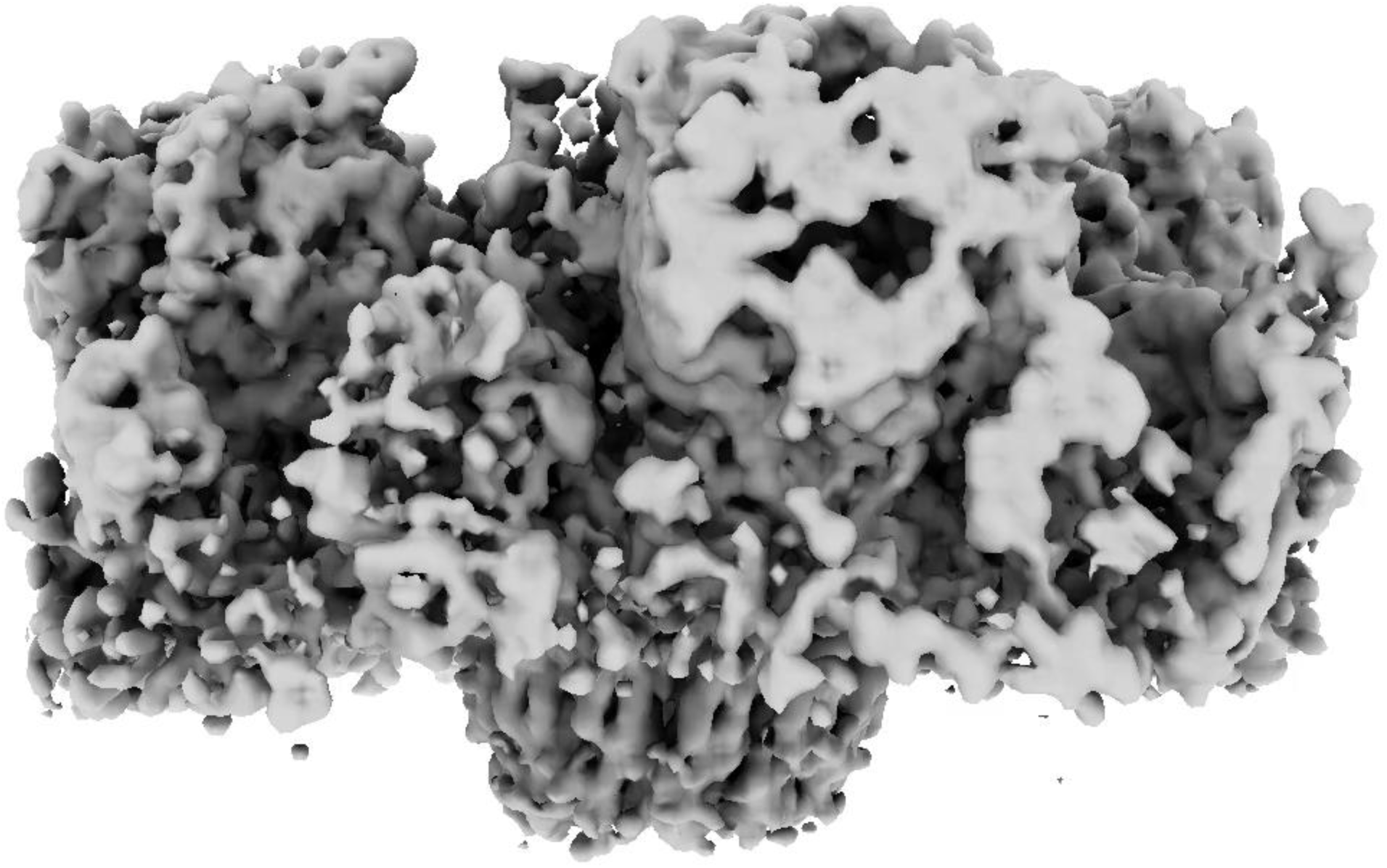
3DFlex volume series of the A-capsid portal vertex, showing slight sway of the basket relative to the capsid shell, as well as substantial variability in height.

**Movie S2.**
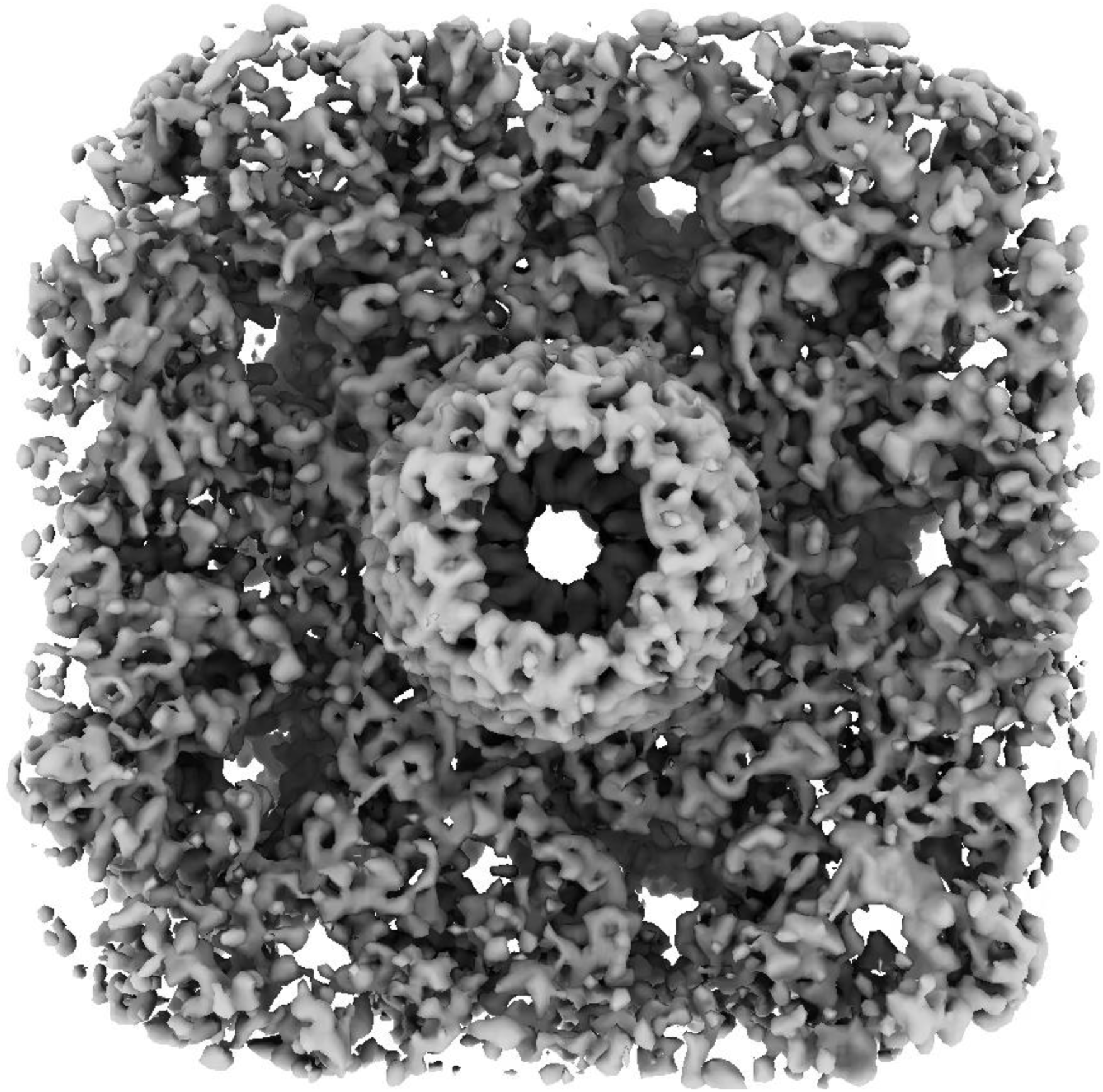
3DFlex volume series of the A-capsid floor at the portal vertex, showing rotation of portal relative to the capsid shell.

**Movie S3.**
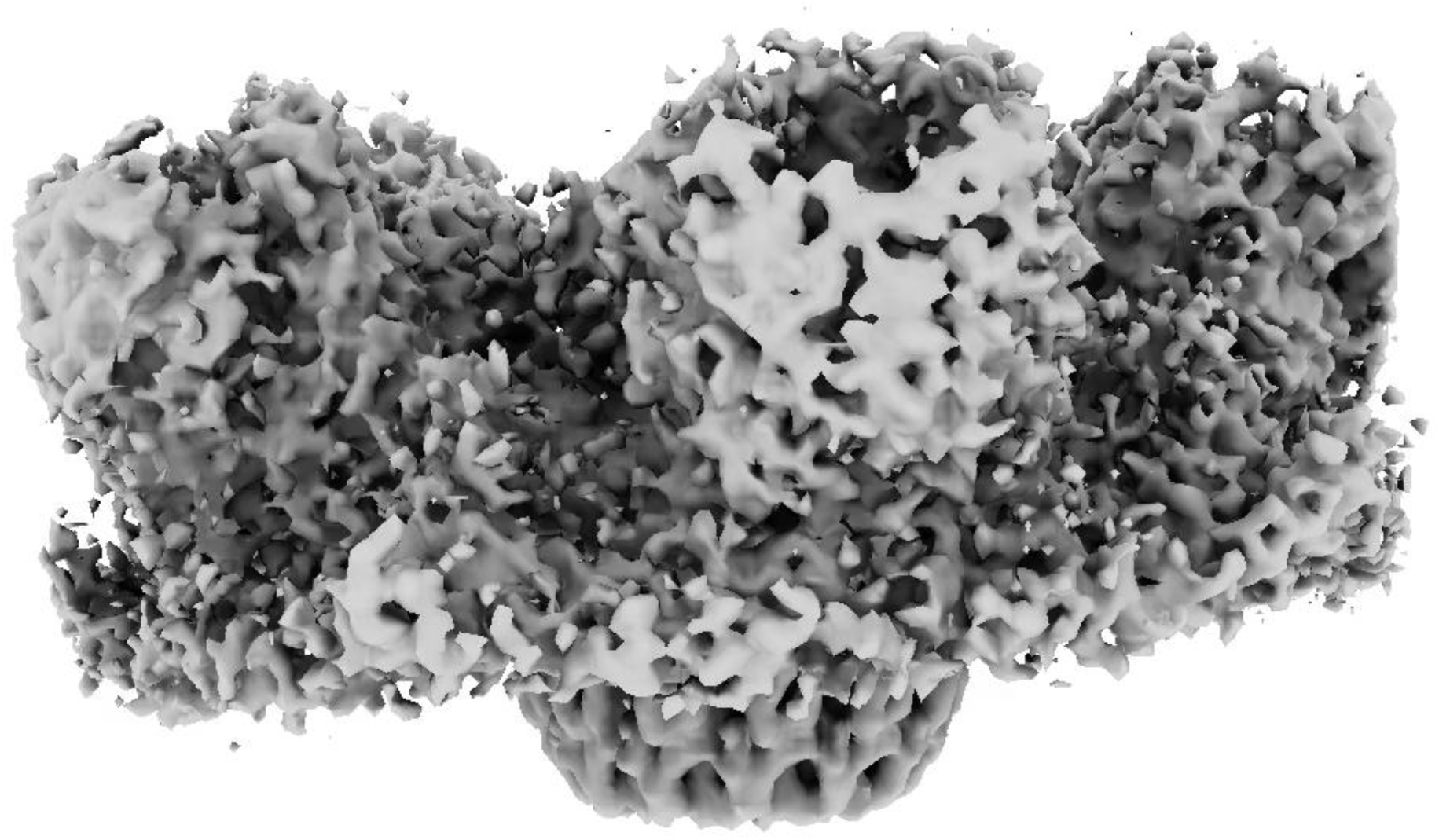
3DFlex volume series of the B-capsid portal vertex, showing sway of the basket relative to the capsid shell.

**Movie S4.**
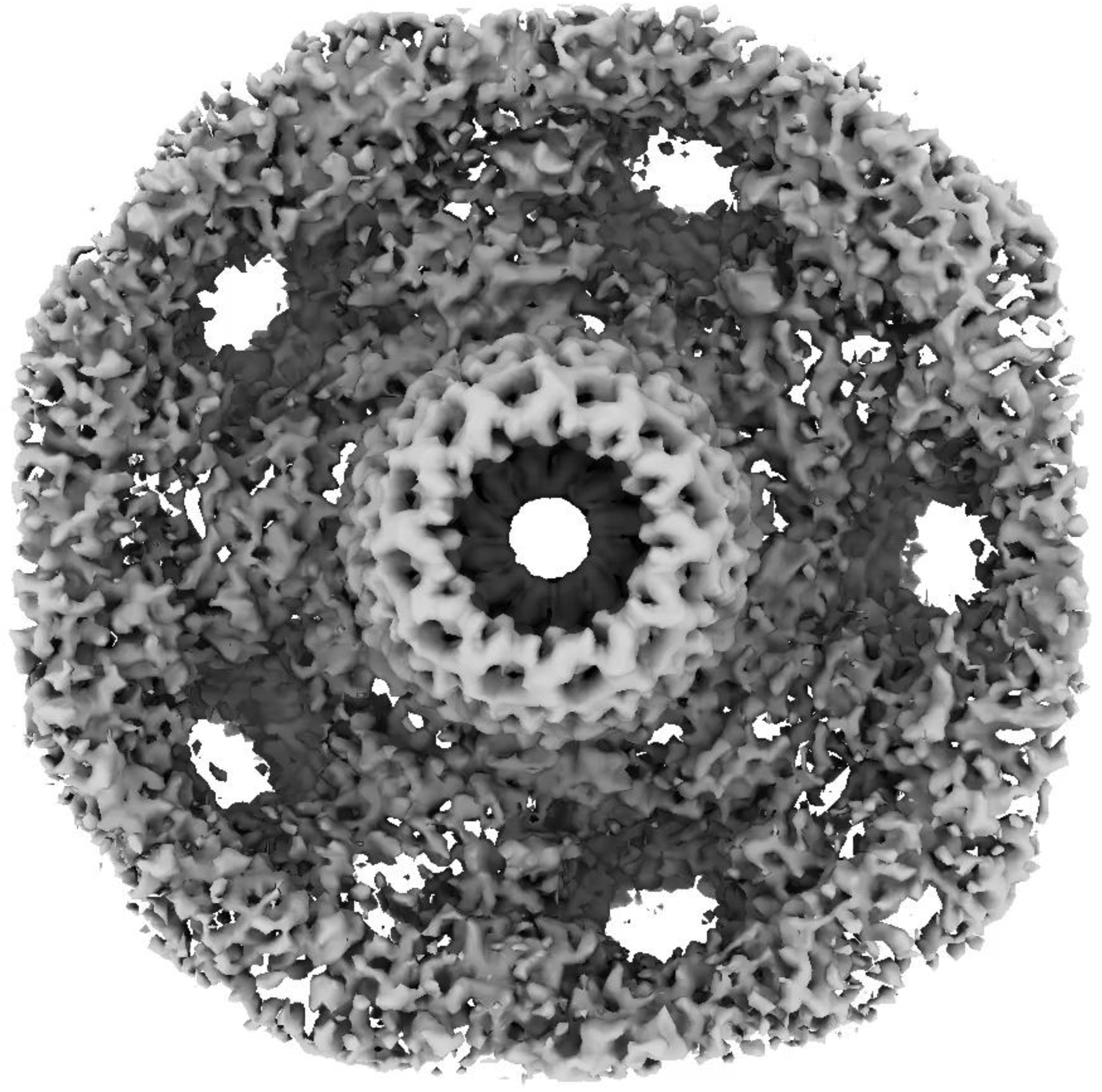
3DFlex volume series of the B-capsid floor at the portal vertex, showing rotation of portal relative to the capsid shell.

**Movie S5.**
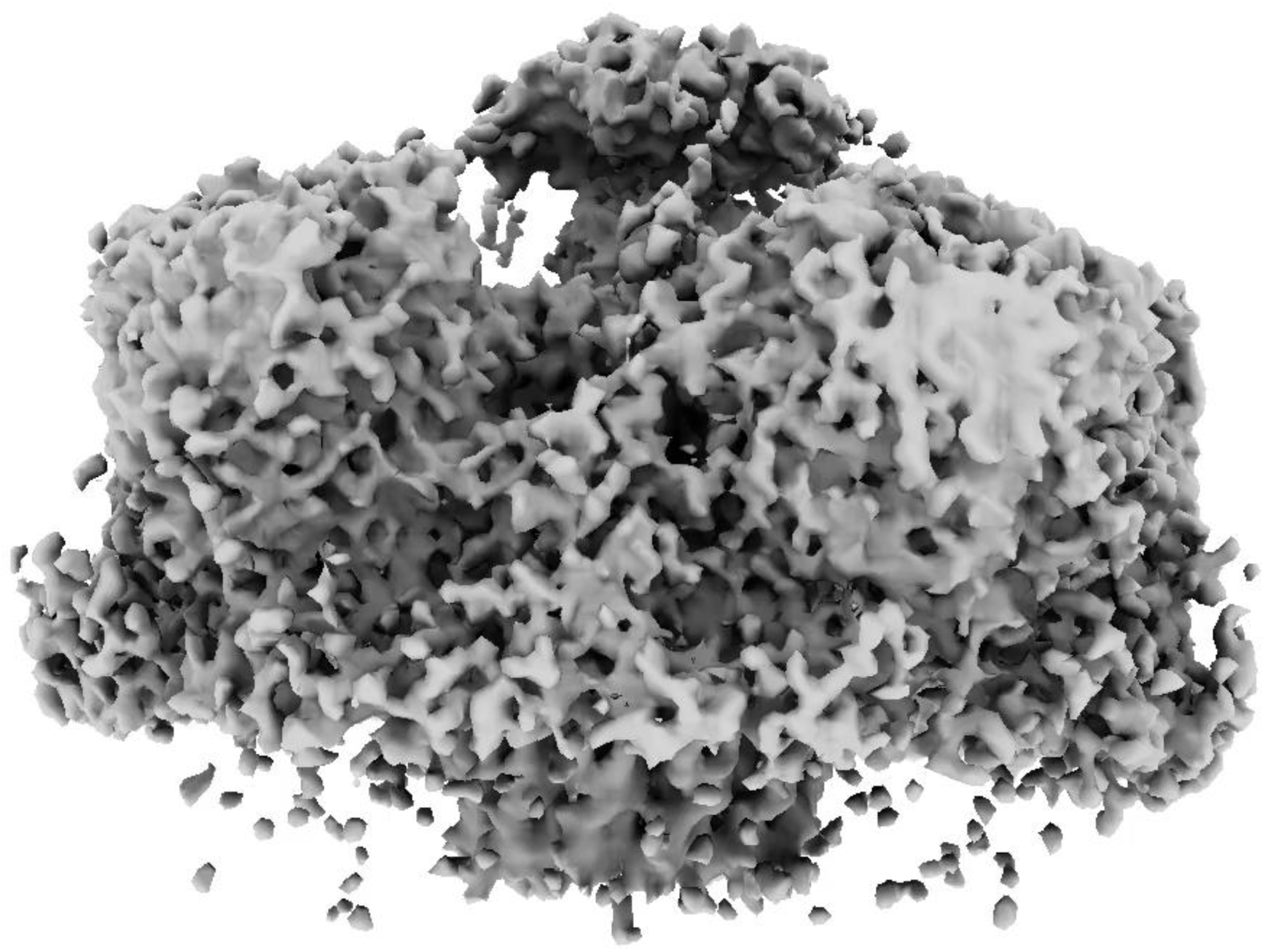
3DFlex volume series of the C-capsid portal vertex, showing sway of the basket relative to the capsid shell.

**Movie S6.**
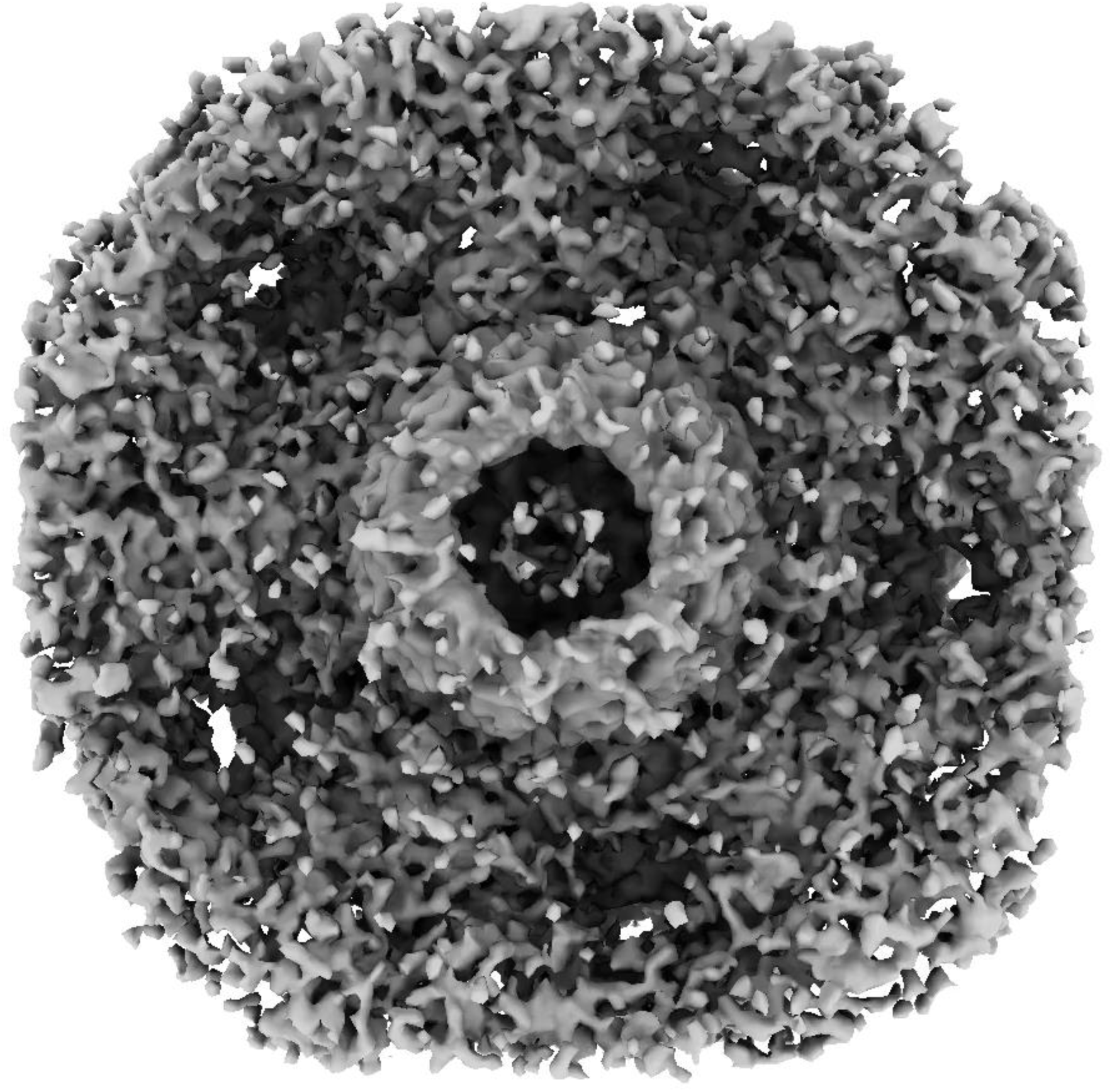
3DFlex volume series of the C-capsid floor at the portal vertex, showing rotation of portal relative to the capsid shell.

**Movie S7.**
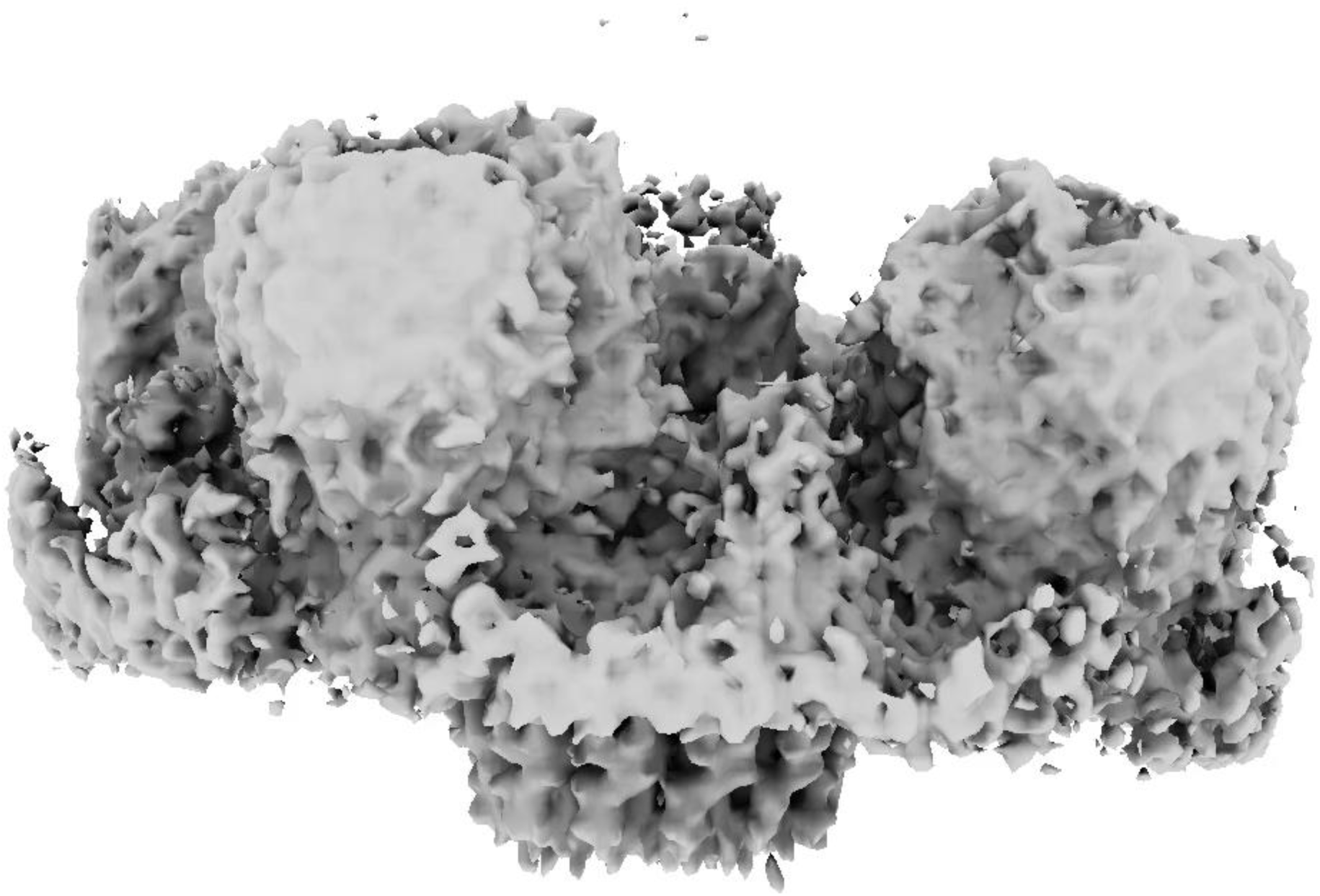
3DFlex volume series of the D-capsid portal vertex, showing sway of the basket relative to the capsid shell.

**Movie S8.**
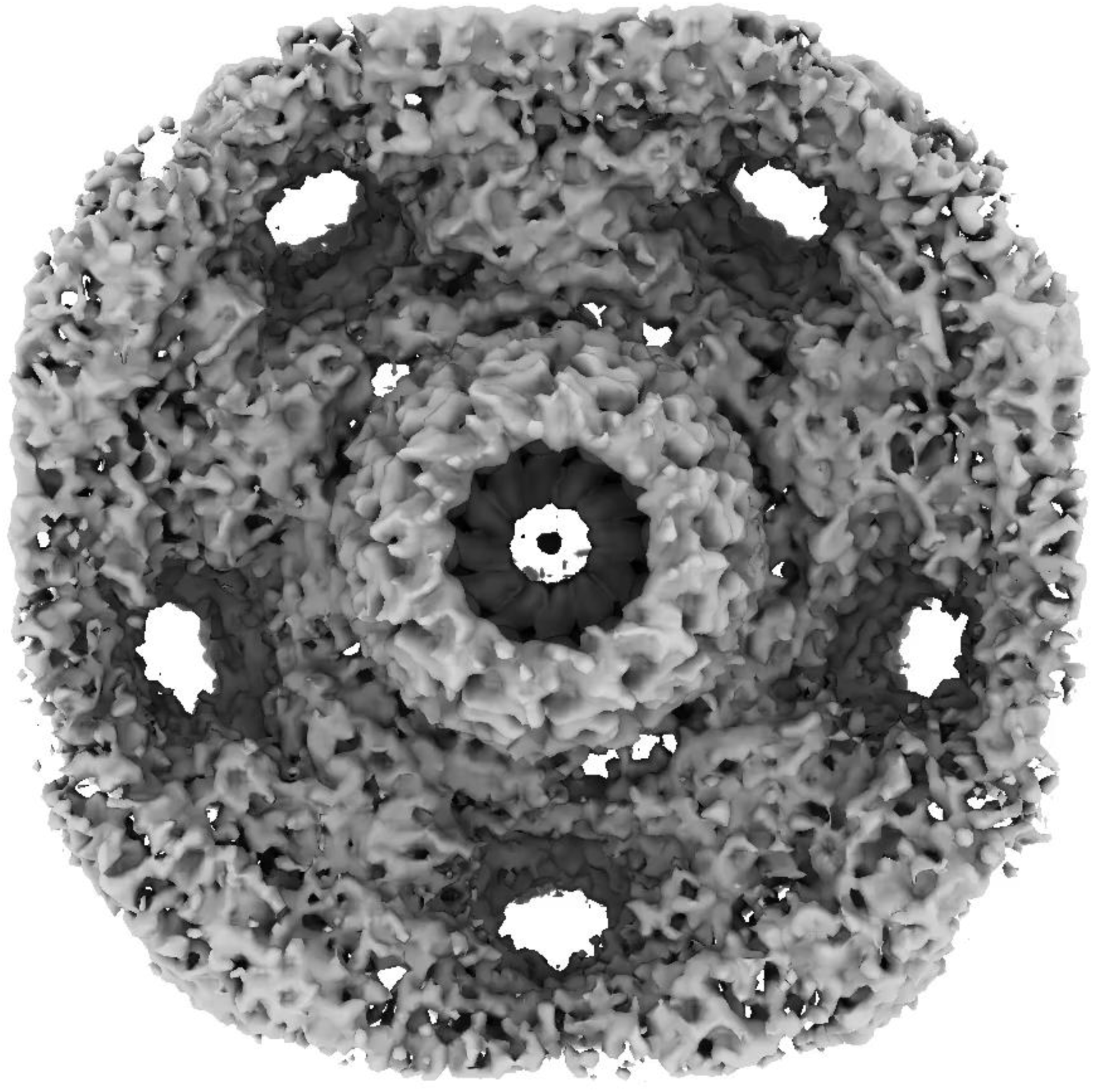
3DFlex volume series of the D-capsid floor at the portal vertex, showing rotation of portal relative to the capsid shell.

